# Systemic delivery of murine SOD2 mRNA to experimental abdominal aortic aneurysm mitigates expansion and rupture

**DOI:** 10.1101/2024.06.17.599454

**Authors:** Huimin Yan, Ying Hu, Yang Lyu, Antonina Akk, Angela C. Hirbe, Samuel A. Wickline, Hua Pan, Elisha D.O. Roberson, Christine T.N. Pham

## Abstract

**Background:** Oxidative stress is implicated in the pathogenesis and progression of abdominal aortic aneurysm (AAA). Antioxidant delivery as a therapeutic for AAA is of substantial interest although clinical translation of antioxidant therapy has met with significant challenges due to limitations in achieving sufficient antioxidant levels at the site of AAA. We posit that nanoparticle-based approaches hold promise to overcome challenges associated with systemic administration of antioxidants.

**Methods:** We employed a peptide-based nanoplatform to overexpress a key modulator of oxidative stress, superoxide dismutase 2 (SOD2). The efficacy of systemic delivery of SOD2 mRNA as a nanotherapeutic agent was studied in two different murine AAA models. Unbiased mass spectrometry-enabled proteomics and high-dimensional bioinformatics were used to examine pathways modulated by SOD2 overexpression.

**Results:** The murine SOD2 mRNA sequence was mixed with p5RHH, an amphipathic peptide capable of delivering nucleic acids *in vivo* to form self-assembled nanoparticles of ∼55 nm in diameter. We further demonstrated that the nanoparticle was stable and functional up to four weeks following self-assembly when coated with hyaluronic acid. Delivery of SOD2 mRNA mitigated the expansion of small AAA and largely prevented rupture. Mitigation of AAA was accompanied by enhanced SOD2 protein expression in aortic wall tissue. Concomitant suppression of nitric oxide, inducible nitric oxide synthase expression, and cell death was observed. Proteomic profiling of AAA tissues suggests that SOD2 overexpression augments levels of microRNAs that regulate vascular inflammation and cell apoptosis, inhibits platelet activation/aggregation, and downregulates mitogen-activated protein kinase signaling. Gene set enrichment analysis shows that SOD2 mRNA delivery is associated with activation of oxidative phosphorylation, lipid metabolism, respiratory electron transportation, and tricarboxylic acid cycle pathways.

**Conclusions:** These results confirm that SOD2 is key modulator of oxidative stress in AAA. This nanotherapeutic mRNA delivery approach may find translational application in the medical management of small AAA and the prevention of AAA rupture.

## Introduction

AAA is an inflammatory disease process characterized by transmural infiltration of the aortic wall with every type of leukocytes^1, 2^ that release matrix metalloproteinases (MMPs) and pro-inflammatory cytokines, such as tumor necrosis factor alpha (TNF-α), interleukin-1β (IL-1β), IL-6, IL-12, IL-23, interferon gamma (IFN-γ), leading to the degradation of extracellular matrix and acceleration of aneurysmal expansion.^1, 3–6^ In addition to matrix-degrading proteases and inflammatory cytokines, infiltrating inflammatory cells can produce large amounts of reactive oxygen species (ROS) that amplify oxidative stress. Studies in human patients and animal models of AAA suggest that oxidative stress contributes to the production of ROS. For example, upregulation of nitric oxide (NO) induces the expression of MMPs in aortic endothelial cells, which exacerbates experimental AAA.^7–9^ Zhang *et al.* showed that inducible nitric oxide synthase (iNOS) is present in human AAA and promotes tissue and cellular injury.^10^ Indeed, inhibition of iNOS with the selective inhibitor aminoguanidine limits experimental AAA.^11^ Despite promising experimental studies pointing to the pathogenic role of ROS, clinical translation has met with significant challenges due to limitations in achieving sufficient levels of antioxidant therapeutics locally at the site of AAA following systemic delivery. Indeed, no single antioxidant therapy has proven successful in clinical trials.

Conversely, enzymes that can eliminate ROS such as catalase, superoxide dismutase (SOD), and glutathione reductase are relatively deficient in AAA tissues,^12^ resulting in an imbalance of oxidants and antioxidants and an excess of ROS production that perpetuate the inflammatory cycle to activate MMPs and induce cellular apoptosis. Yet, few studies have explored the overexpression of antioxidants as a potential therapy. Parastatidis *et al*. showed that overexpression of catalase, a peroxisomal H_2_O_2_ scavenging enzyme, *via* parenteral administration of catalase or by using a genetically engineered murine model, protected mice from CaCl_2_-induced AAA.^13^ In both approaches, catalase was infused either continuously through a pump prior to CaCl_2_ application or was overexpressed from birth by genetic engineering, neither of which may be easily translated into clinical practice.

To overcome potential challenges associated with the practical overexpression of antioxidants, we turned to a nanoparticle (NP)-based approach because NP delivery strategies are postulated to increase therapeutic payloads at local tissue sites while potentially decreasing the risks of adverse events due to systemic administration. Herein, we employed a peptide-nucleic acid nanoplatform^14^ to effect efficient overexpression of SOD2 in aortic tissues. SOD2, one of three types of cellular SODs and the resident mitochondrial form of SOD, also is known as manganese SOD (Mn-SOD2). Because it plays a key role in modulating oxidative stress to protect against mitochondrial damage,^15,16^ we postulated that SOD2 overexpression in aortic aneurysms might forestall the oxidative damage that contributes to aneurysmal expansion and rupture.

## Methods

### Data Availability

The data supporting findings are available from the corresponding authors upon reasonable request. Detailed descriptions of experimental methods, materials, and statistical analysis are presented in the Supplemental Material. Please see the Major Resources Table in the Supplemental Material.

## Results

### Oxidative stress in elastase-induced AAA

We first used the well-established elastase-induced AAA model in which transient porcine elastase perfusion of the infrarenal abdominal aorta on day 0 leads to aneurysmal dilatation at day 14.^19^ AAA is defined as an increase in the aortic diameter (AD) of more than 100% over the pre-elastase perfusion measurements.^19^ Elastase perfusion led to an immediate increase in AD of ∼70% (Figure 1A).^20^ WT C57BL/6 mice uniformly developed AAA on day 14 (mean increase in AD = 154 ± 4% or 0.78 ± 0.02 mm, n = 24) (Figure 1A-B). AAA was accompanied by fragmentation of the elastic fibers (Figure 1C) and an increase in MMP activity (Figure 1D).

**Figure 1.**
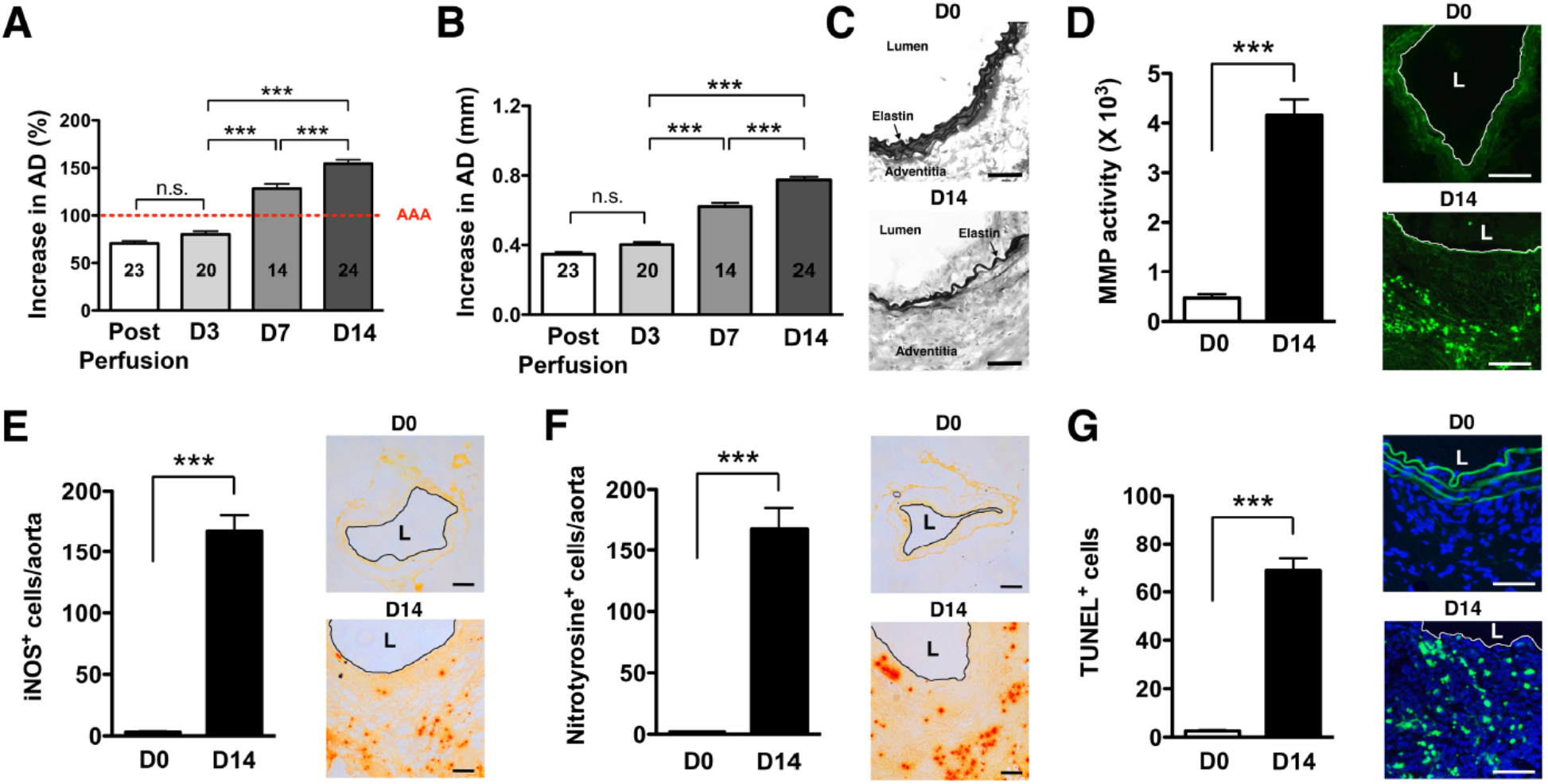
Oxidative stress in elastase-induced AAA. (A) Mice were transiently perfused with elastase on day 0. Aortic diameter (AD) increase is expressed in % or mm. (A) There was an immediate increase in aortic diameter (AD) of *∼*70% immediately post-perfusion. AAA was defined as an increase in AD of greater than 100% compared with the AD measured prior to elastase perfusion. (B) Increase in AD, measured on day 14 as mm increase. Aortic sections from day 14 were examined for elastic fiber integrity with VVG staining (C), in situ MMP activity (D), iNOS by immunohistochemistry (E), NO by nitrotyrosine levels (F) and apoptotic cells by TUNEL staining (G). Values represent mean ± SEM derived from 4-6 non-overlapping fields per aortic section and 3-5 sections per aorta, n = 5-6 aortas per treatment. ***P < 0.001, n.s. not significant. L = lumen. Scale bars = 100 μm (D, E-F), 50 μm (C, G).

Signs of oxidative stress included marked induction of iNOS (Figure 1E), elevated levels of nitrotyrosine (a maker of ROS/NO production, Figure 1F), and an abundance of apoptotic/TUNEL^+^ cells (Figure 1G) on day 14.

### Nanoparticle synthesis and characterization

We have recently shown successful *ex vivo* delivery of long mRNA sequences (> 1,000 nucleotides, nt) to cartilage explants using the p5RHH peptide-based platform.^21^ Building on this early success, we next generated p5RHH-SOD2 mRNA NP. The SOD2 mRNA sequence was produced commercially and contained the appropriate endcaps, poly-A tail, and nucleotide base modifications (∼1,000 nt) to enhance translation. The sequence also contains a mitochondrial localizing peptide component. The p5RHH-SOD2 mRNA NP was prepared by mixing 1 μg of SOD2 mRNA and 10 μmol of p5RHH peptide at 37°C for 40 min to form a self-assembled stable and uniform NP of ∼50-55 nm in diameter by TEM (Figure 2A). The NP was further functionalized with a coating of hyaluronic acid (HA), with the goal of enhancing cellular uptake via CD44 binding.^22^ The highly negative HA coating lowered the NP’s zeta potential to ∼ −30 mV, as previously shown.^22^ We showed that the HA-coated p5RHH-SOD2 mRNA NP (HA-SOD2 mRNA NP) was readily taken up by bone-marrow-derived macrophages (Suppl. Figure 1 and also as previously demonstrated)^22^ and the overexpression of SOD2 was readily detected in mitochondria of RAW 264.7 following *in vitro* transfection, as evidenced by colocalization with Mito tracker red (Figure 2B). The transfected RAW 264.7 cells were lysed, fractionated on SDS-PAGE, and probed with a SOD2-specific antibody to further confirm overexpression (Figure 2C).

**Figure 2.**
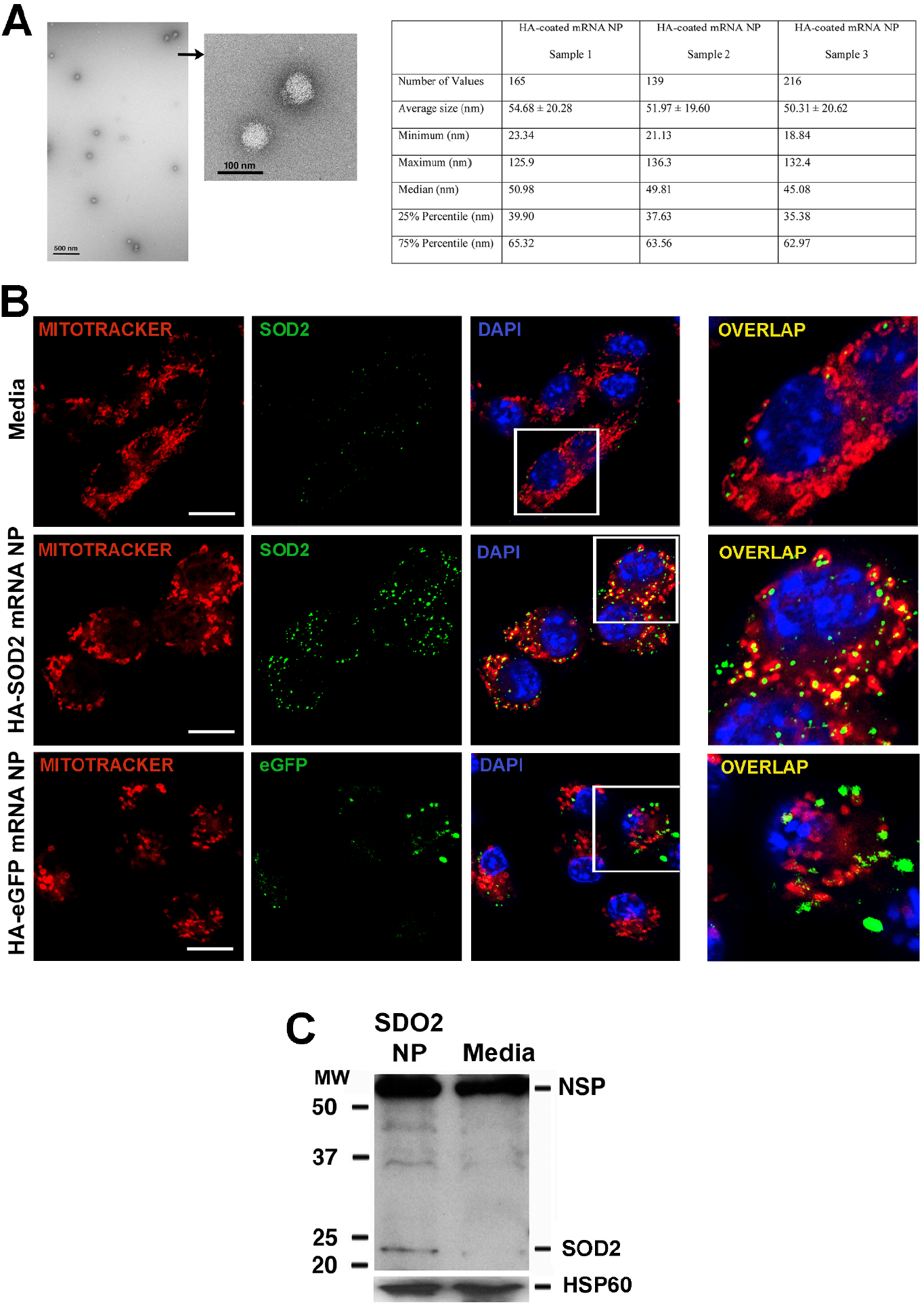
Characterization/uptake of HA-SOD2 mRNA NP and *in vitro* expression of SOD2. NP was prepared by mixing SOD2 mRNA (1 µg) with p5RHH (10 µmol) at 37°C for 40 min. 5 µl of HA was added to the self-assembled NP and placed on ice for 5 min. (A) TEM of HA-SOD2 mRNA NP and sizing from 3 separate NP samples prepared simultaneously. Scale bars = 500 nm; high magnification 100 nm. (B) RAW 264.7 cells were seeded in 8-well Nunc™ Lab-Tek™ II Chamber Slide™ Glass slide System and transfected with HA-SOD2 mRNA NP or HA-eGFP mRNA NP (as irrelevant mRNA control) for 5 hours then the expression of SOD2 and eGFP were detected at 48 hours. The images were captured by a ZEISS LSM 880 confocal laser scanning microscope. Scale bars = 10 μm. (C) RAW 264.7 cells were kept in media or transfected with HA-SOD2 mRNA NP for 5 hours then harvested at 48 hours. Mitochondria were isolated according to manufacturer’s directions, fractionated on SDS-PAGE, and blotted for SOD2. HSP60 served as control for protein loading; NSP = non-specific band.

### Nanoparticle-based delivery of SOD2 mRNA in AAA mitigation

We administered freshly prepared fluorescein-labeled HA-SOD2 mRNA NP i.v. to mice on day 9 post-elastase perfusion. We observed NP localizing to the elastase-perfused abdominal aorta at 7 and 24 hours after injection (Suppl. Figure 2A) with low level sequestration in the liver and minimal accumulation in other major organs (Suppl. Figure 2B). These results confirmed that the incorporation of mRNA instead of small interfering (si)RNA did not significantly affect the localization or biodistribution of HA-p5RHH-based NP in the elastase-induced AAA model.^22^

Next, the fluorescein-labeled HA-SOD2 mRNA NP was administered i.v. to mice on days 5, 8, and 11 post-elastase perfusion. On day 9, 24 hours after i.v. administration NP could be observed colocalizing with MOMA-2^+^ cells in the aortic wall adventitial layer (Suppl. Figure 2C). It should be noted that it is unlikely fluorescein-labeled HA-SOD2 mRNA NP reached the aortic wall tissue mainly *via* cellular hitchhiking as we only detected a small percentage of monocytes (6-8%) with internalized NP in the circulation following i.v. administration (Suppl. Figure 3).

Mice were then perfused with elastase on day 0 and administered HBSS or HA-SOD2 mRNA NP i.v. on days 5, 8 and 11 post-elastase perfusion (mRNA = 1 μg per treatment) (Figure 3A). NP-based mRNA delivery led to enhanced SOD2 expression while suppressing nitrotyrosine levels in aortic wall tissue (Figure 3B-C). HBSS had no effect on SOD2 or nitrotyrosine expression (Figure 3B-C). SOD2 mRNA delivery significantly mitigated AAA progression (AD = 0.57 ± 0.02 mm in HA-SOD2 mRNA NP vs AD = 0.79 ± 0.02 mm in HBSS, P < 0.001) (Figure 3D-E). Mitigation of AAA was accompanied by marked decreases in iNOS^+^, MMP^+^ and TUNEL^+^ cells (Figure 3F-H).

**Figure 3.**
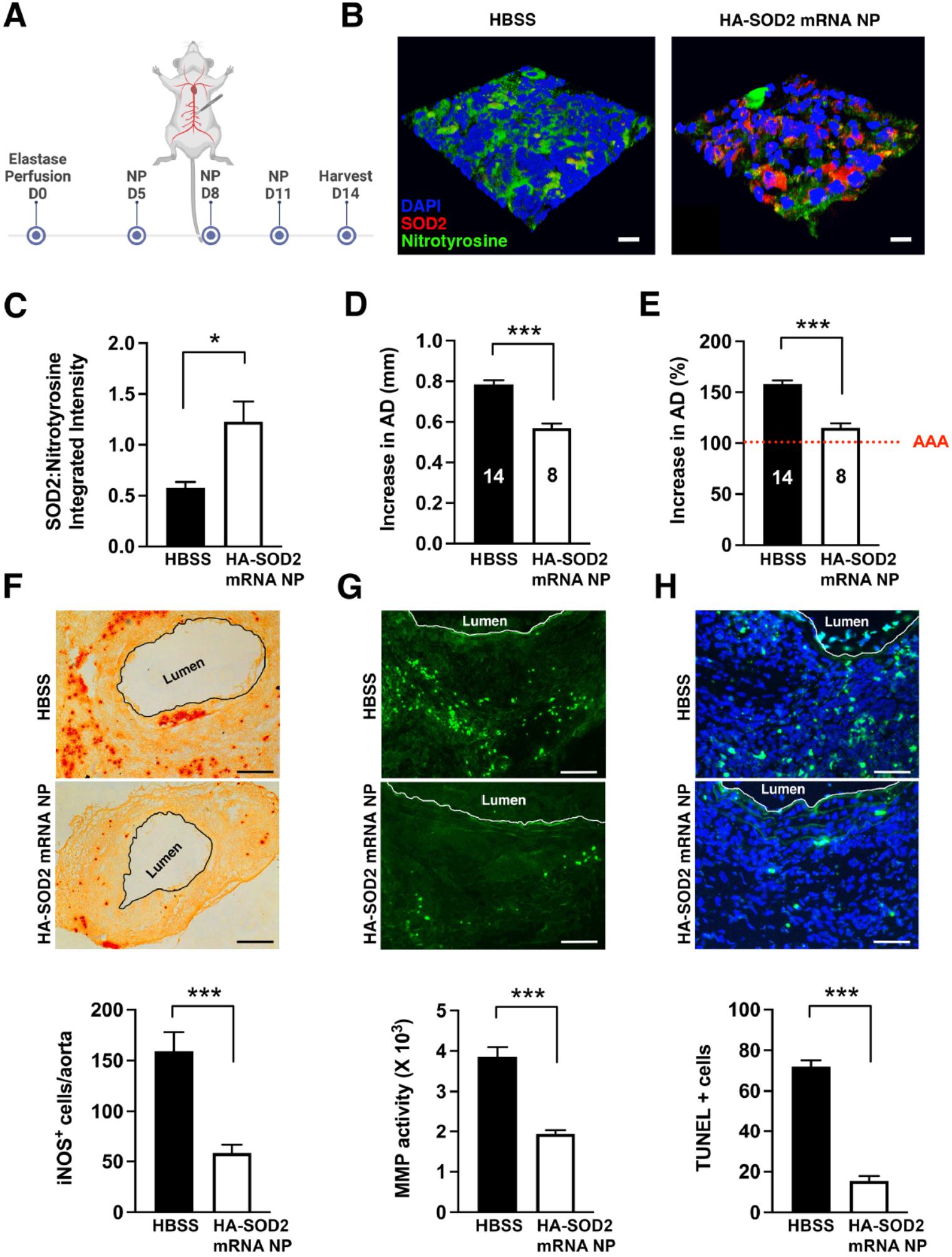
HA-SOD2 mRNA NP in elastase-perfusion model of AAA. (A) Mice were perfused with elastase on day 0 and administered HBSS or HA-SOD2 mRNA NP i.v. on days 5, 8 and 11 post-elastase perfusion (mRNA = 1 μg per treatment). (B) Confocal microscopy of day 14 AAA tissue in HBSS and HA-SOD2 mRNA NP treated animals. SOD2 (red), nitrotyrosine (green), DAPI (blue). Scale bars = 15 μm. (C) Ratios of SOD2:nitrotyrosine intensity. Day 14 aortic diameter (AD) increase was expressed in mm (D) or % (E). Day 14 iNOS^+^ cells (F), *in situ* MMP activity (G), and TUNEL^+^ cells (H) in AAA tissue were assessed. Values represent mean ± SEM derived from n = 4-6 sections per aorta, n = 4-8 aortas per treatment. Scale bars = 200 μm (F), 100 μm (G), 50 μm (H). *P < 0.05, **P < 0.01, ***P < 0.001, n.s. not significant.

### Long-term stability of HA-SOD2 mRNA NP

To test the storage stability and functionality of the HA-SOD2 mRNA NP, we prepared separate batches of NP and refrigerated them at 4°C for 1 to 4 weeks. The NP was assessed weekly for size, zeta potential, and polydispersity. We observed that the HA-coated NP was remarkably stable over time (Suppl. Figure 4). Following *in vitro* transfection into RAW 264.7 cells with the 4-week-old stored NP, we still observed strong mitochondrial overexpression of SOD2 (Figure 4A). Moreover, the NP was functional after 4 weeks in storage at 4°C, as evidenced by their ability to mitigate AAA as efficiently as freshly prepared NP (Figure 4B-C). We confirmed the suppression of oxidative stress, as shown by decrease in iNOS, MMP, and TUNEL-positivity (Figure 4D-F). Lastly, administration of NP did not affect hematologic parameters or liver/kidney function (Suppl. Figure 5).

**Figure 4.**
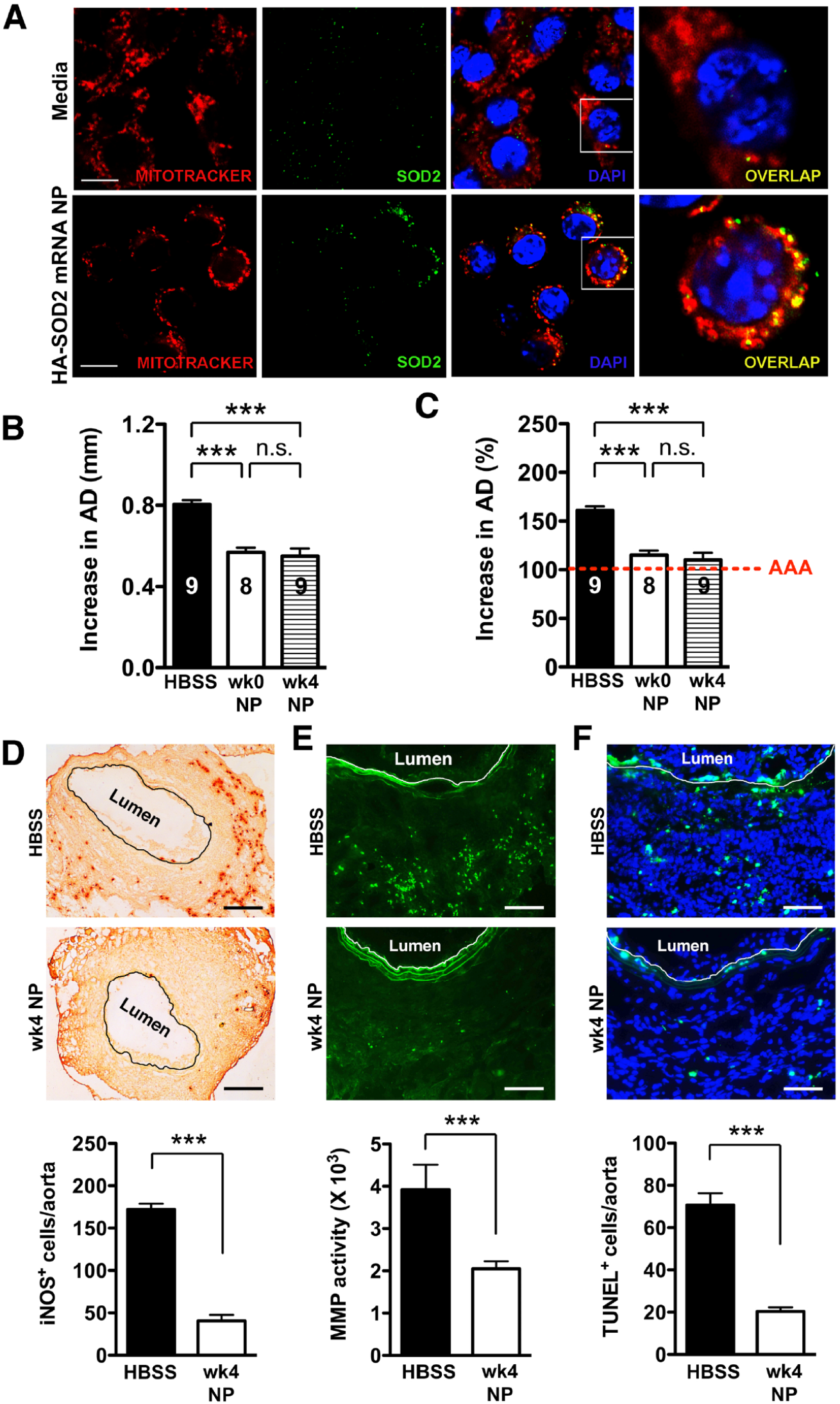
Long-term stability of HA-SOD2 mRNA NP. (A) RAW 264.7 cells were seeded in 8-well Nunc™ Lab-Tek™ II Chamber Slide™ Glass slide System and transfected for 5 hours with HA-SOD2 mRNA NP, which has been stored at 4°C for 4 weeks and the expression of SOD2 was examined at 48 hours. The images were captured by a ZEISS LSM 880 confocal laser scanning microscope. Mice were perfused with elastase and on days 5, 8 and 11 were administered HA SOD2 mRNA NP that was freshly prepared or has been stored at 4°C for 4 weeks. Mice were sacrificed on day 14 and aortic diameter (AD) was measured. Increase in AD was expressed in mm (B) or % (C). Day 14 iNOS^+^ cells (D), in situ MMP activity (E), and TUNEL^+^ cells (F) in AAA tissue were assessed. Values represent mean ± SEM derived from n = 4-6 sections per aorta, n = 4-8 aortas per treatment. Scale bars = 10 μm (A), 200 μm (D), 100 μm (E), 50 μm (F). ***P < 0.001, n.s. not significant.

### Nanoparticle-based delivery of SOD2 mRNA in the prevention of AAA rupture

To examine the role of SOD2 in AAA rupture, we turned to a TGF*β*-blockade model of AAA rupture, which was developed by Lareyre and colleagues.^17^ Peri-aortic application of elastase to the abdominal aorta combined with systemic blockade of TGFβ activity led to rapid aortic diameter expansion and rupture in ∼50% of experimental animals within 14 days. As described in our previous study,^22^ this model also exhibits signs of oxidative stress with robust level of iNOS accompanied by marked production of MMPs, aortic tissue remodeling, apoptosis and intraluminal thrombus.

In this model, elastase was applied peri-adventitially to the abdominal aorta followed by systemic administration of TGFβ blocking antibody on days 0, 3, 5, 7, 10, 13 (Figure 5A) leading to marked enlargement of the abdominal aorta (Figure 5B) and ∼55% rupture in our hands, without intervention (Figure 5C). HA-SOD2 mRNA NP administration not only delayed the onset of rupture (until day 11) but significantly protected mice from sudden death (87.5 % survival rate in treated mice, n=20 vs 45 % survival rate in untreated mice, n=16, P < 0.01) (Figure 5C). In addition, HA-SOD2 mRNA NP treatment also mitigated the progression of AAA growth in 80% of the surviving animals (11 out of 14 surviving mice, Figure 5D-E). Mitigation of AAA growth was accompanied by significant reduction of iNOS, TUNEL+ cells and *in situ* MMP activity (Figure 5F-H). Lastly, overexpression of SOD2 markedly suppressed the level of nitrotyrosine, a marker of oxidative stress-related pathologies (Figure 6).

**Figure 5.**
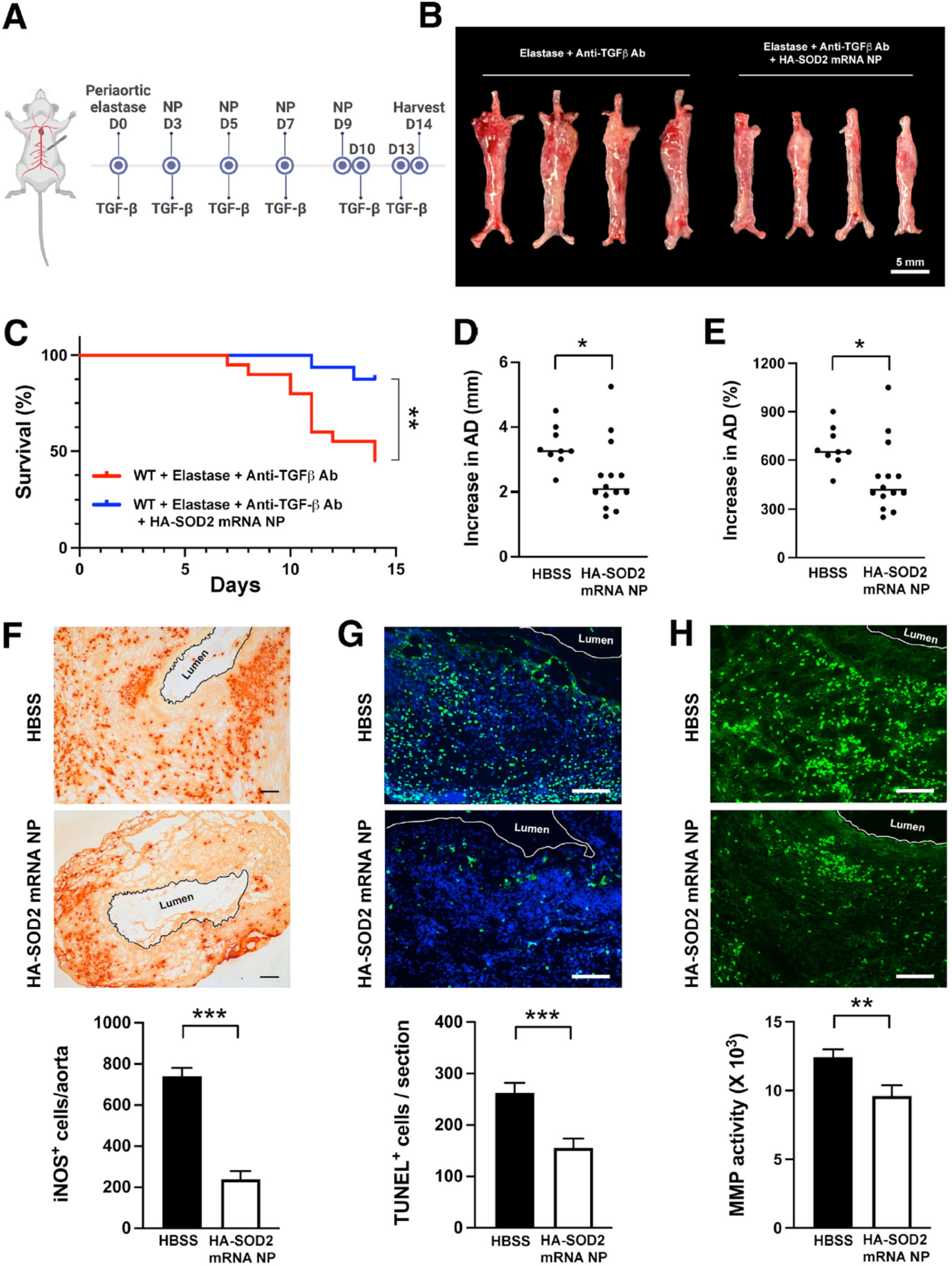
HA-SOD2 mRNA NP in TGFβ blockade model of AAA rupture. Periaortic elastase was applied on day 0; TGFβ antagonist and NP were administered according to the schedule shown in (A). (B) Representative macroscopic images of aortas on day 14. Scale bar = 5 mm. (C) Survival curves following the different treatment regimens. The surviving mice were sacrificed on day 14 and aortic diameter (AD) was measured. Increase in AD was expressed in mm (D) or % (E). Day 14 iNOS+ cells (F), TUNEL+ cells (G), and *in situ* MMP activity (H) in AAA tissue were assessed. Values represent mean ± SEM derived from n = 4–6 sections per aorta, n = 4–8 aortas per treatment. Scale bars = 100 μm (F, G, H). *P < 0.05, ***P < 0.001, n.s. not significant.

**Figure 6.**
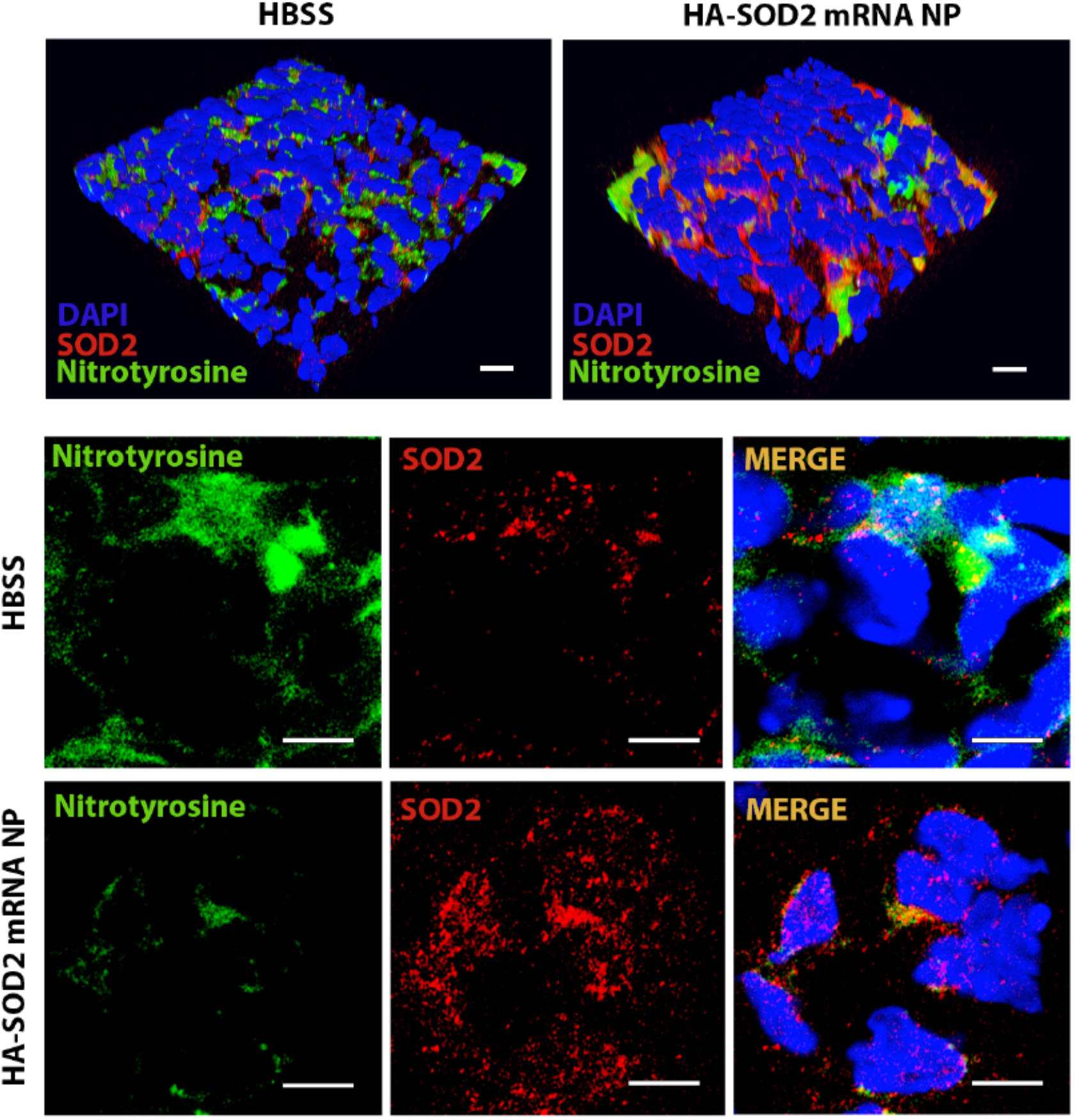
SOD2 overexpression in vivo following HA-SOD2 mRNA NP administration. Aortic sections from day 14 TGFβ blockade model of AAA rupture was examined for NO (nitrotyrosine, green) and SOD2 (red) levels. Scale bars = 100 μm (upper panel), 50 μm (lower panel).

### Proteomic profiling of TGFβ blockade model of AAA rupture

To further explore and identify proteins and molecular pathways that accompanied SOD2 overexpression, we turned to unbiased mass spectrometry-enabled proteomics and high-dimensional bioinformatics to provide the complete proteome profiling of HA-SOD2 mRNA NP-treated mice compared to non-treated animals. We analyzed non-imputed data to arrive at the complete proteome profiles of 4 HA-SOD2 mRNA NP-treated (T1-4) and 4 non-treated abdominal aortas (NT1-4) of day 14 surviving animals from the TGF*β*-blockade model. Principal Component Analysis (PCA) was conducted to assess sample distribution and distinguish the non-treated vs. treated samples based on their protein abundance/expression profiles (Figure 7A). A heatmap of differentially expressed proteins is shown in Figure 7B. The upper split revealed that NT1 & NT2 have high detection while NT3 and NT4 have medium detection (Figure 7B); treated T1-4 have low detection, (Figure 7B). Compared with treated aortas, which have an average AD of 3.25-3.30 mm, NT1 and NT2 have an average AD of 4.00-4.50 mm and exhibited localized hemorrhage, suggesting impending rupture. Thus, the high detection of Plex (pleckstrin), Trpc6 (transient receptor potential channel 6), Rasa3 (Ras p21 protein activator 3), and Alox12 (arachidonate 12-lipoxygenase) may portend risk of arterial aneurysm rupture while overexpression of SOD2 consistently downregulated these pathways, potentially protecting the animals from aneurysm rupture and sudden death. Subsequent analysis of significantly enriched pathways derived from AAA proteomes revealed that HA-SOD2 mRNA NP treatment modulated cellular response to stress, potentially through the regulation of microRNA (miR)17-5p, which inhibits apoptosis,^23, 24^ and miR 181a-5p, which regulates vascular inflammation/endothelial cell senescence *via* STAT3 signaling pathway (Figure 7C-D).^25, 26^ Furthermore, HA-SOD2 mRNA NP treatment inhibited pathways involved in platelet activation/aggregation (Csk, Lyn, Arrb1, Plek), cell adhesion (Isg15, Lyn), and MAPK1/3 signaling (Rasa3, Csk, Arrb1) (Figure 7C-D). An expanded list of the pathways modulated by SOD2 overexpression can be found in Suppl. Figure 6.

**Figure 7.**
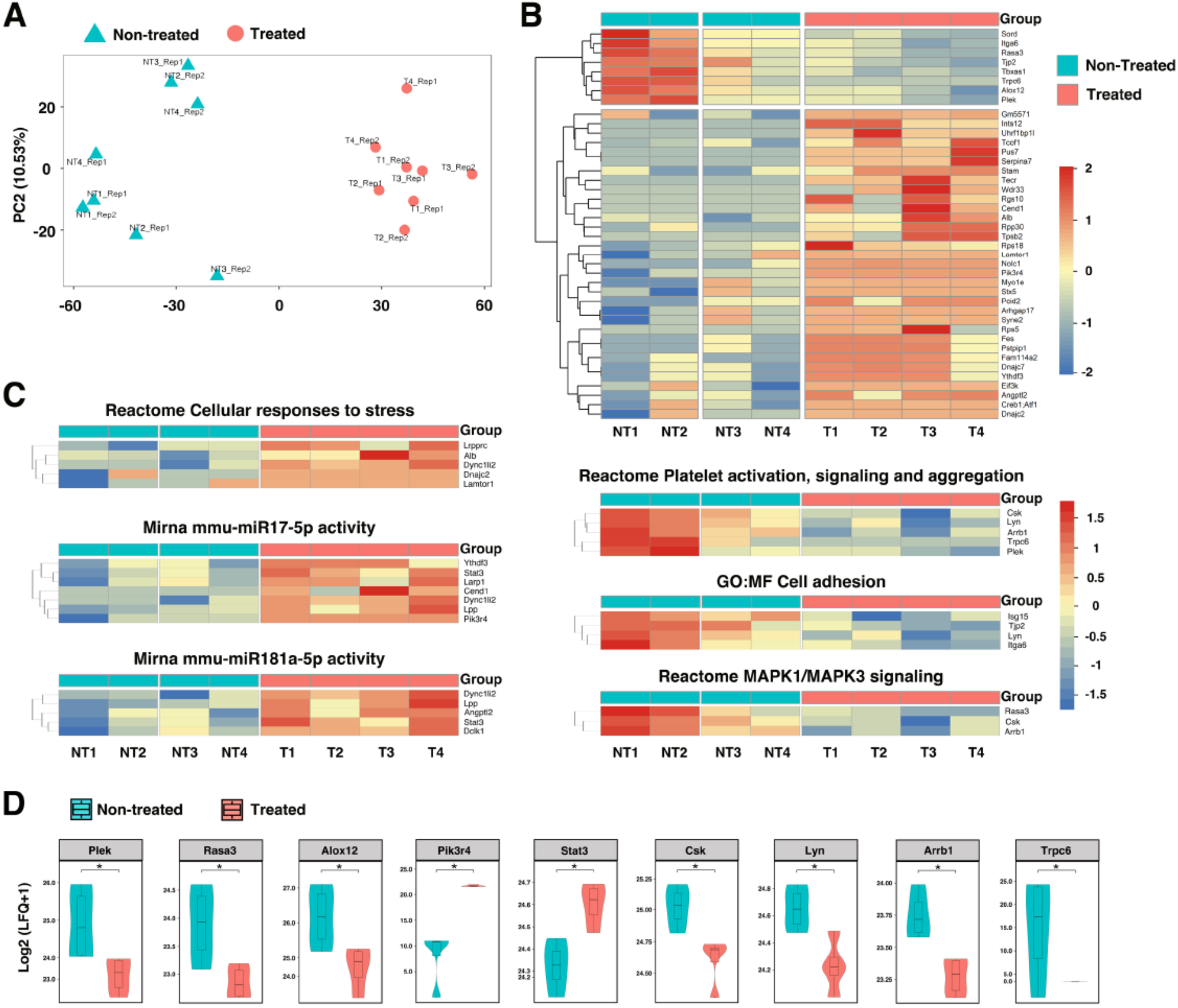
Proteomic profiling in TGFβ blockade model of AAA following HA-coated p5RHH-SOD2 mRNA NP administration. Aortic tissue from non-treated (NT=4) and HA-SOD2 mRNA NP-treated (T=4) were subjected to unbiased mass spectrometry-enabled proteomics and high-dimensional bioinformatics. (A) PCA plot distinguished the non-treated and treated samples based on gene expression profiles. (B) Hierarchical cluster analysis showing relative abundances of proteins in non-treated (NT) and HA-SOD2 mRNA NP-treated (T) aortas. (C) Heatmaps of significantly enriched pathways. (D) Enhancement of key protein components following HA-SOD2 mRNA NP treatment. *P < 0.001.

Because the formation and progression of AAA is usually accompanied by mitochondrial dysfunction and oxidative stress, we sought to further explore molecular pathways and identify effectors participating in mitochondrial biogenesis and the generation of mitochondrial ROS. The majority of ROS (about 90%) is generated during the mitochondrial oxidative phosphorylation process as byproduct. Excessive production of ROS can lead to mitochondrial dysfunction and cell death. Analysis of imputed data revealed significant enrichment in pathways associated with oxidative phosphorylation, lipid metabolism, citric acid tricarboxylic acid (TCA) cycle, and respiratory electron transport in TGF*β*-blockade, AAA rupture model (Figure 8A-B and Suppl. Figure 7). The oxidative phosphorylation system consists of five protein complexes, of which protein complexes I - IV are respiratory chain complexes, including NADH dehydrogenase (NADH-ubiquinone oxidoreductase; complex I), succinate dehydrogenase (complex II), cytochrome c oxidoreductase (cytochrome b and c1; complex III), and cytochrome c oxidase (complex IV).^27^ Enhancement of key protein components of the oxidative phosphorylation system (Ndufa9, Cyc1, Uqcr10, Sdhb/c) were observed after HA-SOD2 mRNA NP treatment (Figure 8C). Proteins in the citric acid TCA/respiratory electron transport systems that were augmented with HA-SOD2 mRNA NP treatment include Etfa/b, Suclg1, Cox6a1 (Figure 8C). Additionally, we observed significant enhancement of Slc25a1, Acadm, Hadha/b, Acaa2 in the fatty acid/lipid metabolism pathways with SOD2 augmentation (Suppl Figure 7). Conversely, we observed decrease in proteins associated with transcription/translation and neutrophil degranulation in animals treated with HA-SOD2 mRNA NP (Figure 8A).

**Figure 8.**
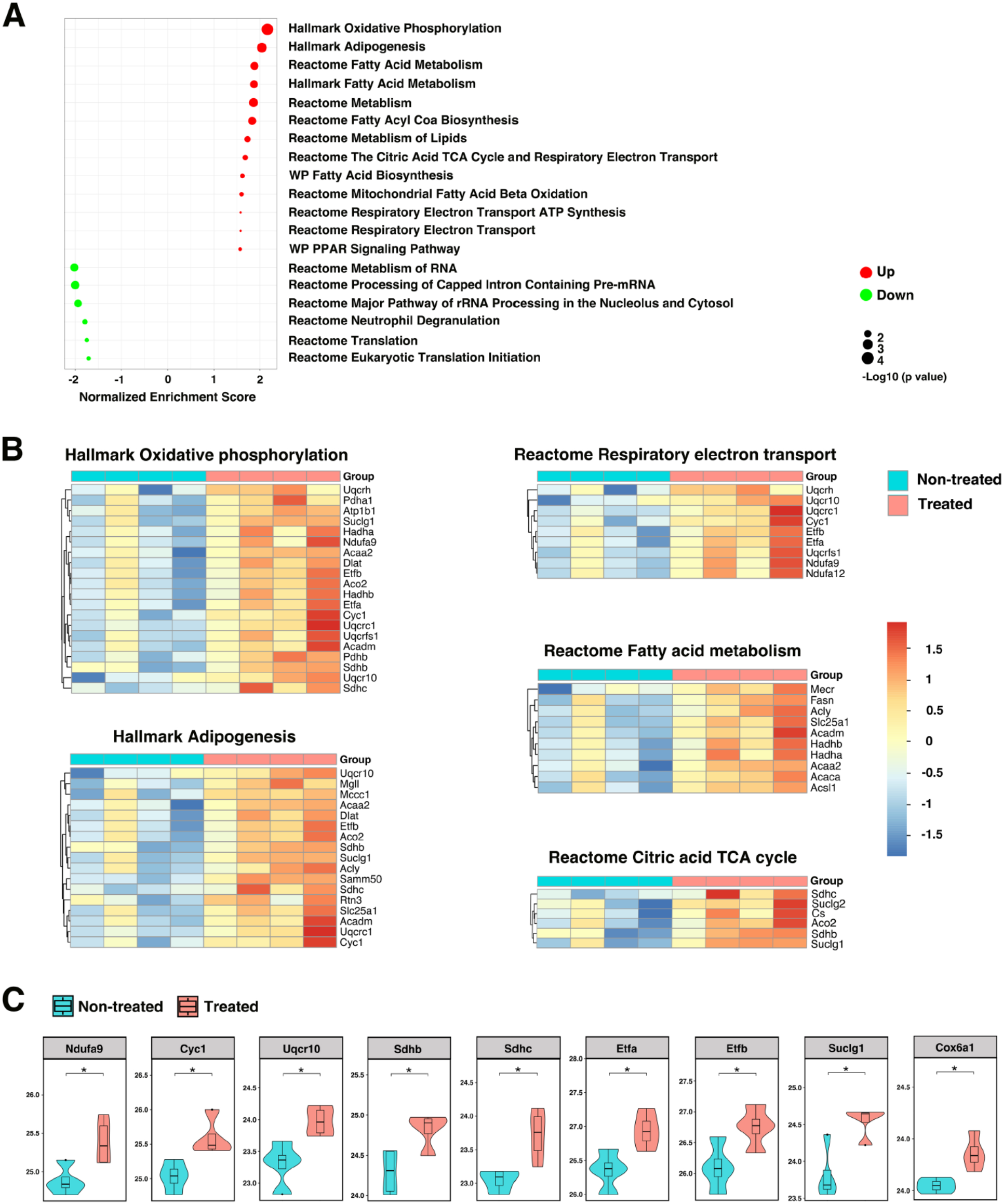
Contribution of SOD2 in the maintenance of mitochondrial redox balance. (A) GSEA revealed significantly enriched pathways in mitochondria following SOD2 augmentation in TGFβ blockade model of AAA. Heatmaps (B) and enhancement of key protein components (C) of pathways that control oxidative-phosphorylation, respiratory electron transport, fatty acid metabolism, and TCA cycle. *P < 0.05.

## Discussion

Nanotechnology-based therapy for AAA treatment is beginning to emerge and promises to overcome challenges presented by systemic delivery of antioxidants.^28^ In this study, we employed a peptide-based NP structure that represents the culmination of a number of specific sequence modifications to the amphipathic cationic peptide, melittin.^29–31^ In this present version of the peptide, called p5RHH, its detrimental pore forming capacity has been substantially attenuated (a 5,700% decrease) while still permitting rapid nucleotide polyplex self-assembly, cellular uptake, prominent endosomal permeabilization and coordinated release of siRNA or mRNA into cytoplasm. ^30, 31^ We also functionalized the peptide-siRNA/mRNA NP with a HA coating to enhance macrophage uptake.^21, 22^

The delivery of mRNA into cells has been attempted using a variety of platforms, including lipid-based nanocarriers. The recent success of COVID-19 vaccines illustrates the potential of mRNA technologies to revolutionize medical care. We have shown previously that delivery of p5RHH-WNT16 mRNA NP to human cartilage explants led to the expression of its downstream effector molecule lubricin, which is important for cartilage protection.^21^ Herein, we demonstrate that the HA-SOD2 mRNA NP effectively suppressed elastase-induced AAA, confirming that the delivery of mRNA using our p5RHH platform also works effectively *in vivo.* Moreover, the HA coating stabilizes the NP, allowing storage for up to 4 weeks post self-assembly. This finding has translational implications as it will allow for storage and transport of the NP agent to the point of care (e.g., the physician’s office) without loss of efficacy. We have also previously shown that the p5RHH-siRNA NP has a favorable safety profile *in vivo*, even after repeated injections.^32, 33^ Likewise, we showed in this study that the delivery of HA-SOD2 mRNA NP did not lead to sustained accumulation of the NP in major organs or affect hematologic parameters or liver/kidney function.

The mitigation of aneurysm *via* overexpression of SOD2 also elucidates the role of this enzyme and ROS in the progression of AAA. The exact contribution of NO to AAA has been difficult to confirm, in part because NO delivery *in vivo* is challenging due to its extremely short half-life.

Likewise, studies using inhibitors to decipher the role of NOS *in vivo* may not be conclusive due to the non-selective nature of these agents. And although SOD2 has been implicated in the pathogenesis of AAA,^34, 35^ overexpression studies have not been attempted. Augmentation of SOD2 using an adenoviral gene delivery approach has been shown to prevent lesion formation in proliferative vascular diseases such as restenosis.^36^ In this restenosis model, adenovirus was introduced into the carotid artery at the time of balloon injury whereas we administered SOD2 mRNA NP on day 5 post-elastase infusion, at a time when the aneurysmal process is already established. Our findings herein suggest that SOD2 overexpression may be a promising therapeutic approach for the treatment of small AAA.

In normal cellular function, the redox status reflects a homeostatic balance between oxidants and antioxidants, preventing excessive generation of ROS that can contribute to cellular damage and the pathophysiology of several diseases. Excessive ROS, particularly within mitochondria, can lead to the destruction of subcellular organelles through various pathways. SOD2 efficiently converts superoxide to the less reactive hydrogen peroxide (H_2_O_2_), which can then freely diffuse across the mitochondrial membrane to undergo breakdown by cytoplasmic peroxidases such as catalase to combat the damaging effects of ROS.^37^ Increasing evidence suggests a key role for mitochondrial damage due to oxidative stress driving AAA pathophysiology.^38^ Moreover, genetic variations in the ROS-generating NADPH oxidase genes have been linked to higher risk of AAA rupture.^39^

Mitochondrial proteins are primary targets for oxidation, causing proteome imbalance that further exacerbates oxidative stress.^40^ Dysfunctional mitochondria due to excessive ROS has been associated with many disorders,^41^ including cardiac diseases,^42^ neurodegenerative diseases,^43^ and aging.^40^ However, the exact role of mitochondrial antioxidants in the progression of AAA remains untested. Herein we showed that SOD2 overexpression in mitochondria mitigates the progression of AAA likely by modulating pathways that control oxidative phosphorylation, respiratory electron transport, fatty acid metabolism, and TCA cycle kinetics. Mitigation of AAA progression and redox modulation is also reflected in the downregulation of platelet activation, MAPK signaling, restriction of vascular inflammation *via* miR-181a-5p, and inhibition of apoptosis *via* miR17-5p and STAT3 signaling.

AAA rupture represents a true medical emergency as it is associated with a mortality rate of ∼81%.^44^ The ability to predict AAA rupture, however, remains an unmet medical need. Currently, the most widely used criterion for prediction of rupture remains the maximum diameter of AAA.^45^ Although other factors such as aortic stiffness have been evaluated,^46^ further validation in larger cohorts is still needed. The rupture of an aneurysm is not a sudden event but rather the culmination of many small occurrences (i.e. microleaks) that lead to serious and potentially fatal consequences. Our proteome profiles revealed enhanced concentrations of Plex, Trpc6, Itg6, Rasa3, Alox12 in non-treated aortas that exhibited localized hemorrhage. Pleckstrin (Plex) is a platelet protein that, upon platelet activation, translocates to the plasma membrane where it is phosphorylated by the protein kinase C (PKC), exerting downstream effects on platelet activation,^47^ and secretion of platelet factors that contribute to thrombus formation^48^ and the risk of rupture.^49^ Trpc6 regulates a multitude of physiologic processes including vascular structural integrity and permeability.^50^ Trpc6 also has a direct role in leukocyte extravasation and endothelial inflammatory response and knockdown of Trpc6 blocks leukocyte transmigration.^51^ Recent data revealed that RASA3, which belongs to the GAP1-family GTPase-activating proteins (GAPs), can act as gatekeeper of T cell activation and adhesion.^52^ Although current evidence supports a pro-inflammatory role of T cells in the progression of AAA,^53^ targeted therapy against T cells in AAA presents significant challenges. Thus, GAPs may offer an alternative T cell targeting strategy. Lipid peroxidation induced by ROS plays a crucial role in various cell death processes, including apoptosis, autophagy, and ferroptosis.^54^ ALOX12 catalyzes the addition of molecular oxygen to arachidonic acid and its activation can enhance lipid ROS accumulation, promoting ferroptosis, which has been implicated in many conditions, including AAA.^55–58^ Taken together, our proteomic analysis identifies a set of proteins that may be explored as markers of impending AAA rupture.

In summary, proteomic profiling and analysis revealed enrichment in pathways related to inflammation/inflammatory responses (i.e., STAT3/MAPK signaling), platelet activity, and translation/transcription, which have been previously demonstrated in other pre-clinical AAA models in addition to oxidative phosphorylation and neutrophil degranulation, pathways that are predominant in human aneurysms.^59^ Moreover, we found that miR181 and miR17 pathways are modulated by SOD2 and may be tracked as biomarkers of disease progression in the circulation. Thus, these findings shed further insights to the contribution of SOD2 in the maintenance of vascular redox balance and reveals several potential therapeutic targets downstream of SOD2 that may be explored as therapeutic targets in the mitigation of AAA progression and rupture. Furthermore, we have established the efficiency of peptide-based delivery of mRNA for AAA treatment that can be extended to the modulation of a multitude of pathways.

Currently there is no selective or molecularly targeted therapy for AAA. Patients may be receiving medications for blood pressure control or lipid lowering that may exert supportive benefit. Open surgical repair or minimally invasive endovascular repair with catheter delivered grafts can be utilized at later stages of the disease to prevent catastrophic rupture and maintain aortic patency. Molecular targeted NP-based nanomedicine offers a promising approach to mitigate AAA progression and rupture prevention, but how this technology will translate to human disease and which time points might be appropriate for initiating SOD2 augmentation remain to be determined in future investigations.

## Acknowledgments

The authors thank Mischa Ahmad for her technical help. Mass Spectrometry analyses were performed by the Mass Spectrometry Technology Access Center at McDonnell Genome Institute (MTAC@MGI) at Washington University School of Medicine. Some figures were created with BioRender.

## Supplemental Material

### Supplemental Methods

Figure S1-S8

Major Resources Table

## Non-standard Abbreviations and Acronyms

AAA: abdominal aortic aneurysm
AD: aortic diameter
HA: hyaluronic acid
IFN: interferon
IL: interleukin
MMP: matrix metalloprotease
NOS: nitric oxide synthase
ROS: reactive oxygen species
iNOS: inducible nitric oxide synthase
SOD: superoxide dismutase
NO: nitric oxide
NP: nanoparticle
TGF*β*: transforming growth factor-*β*
siRNA: small interfering RNA
TNF: tumor necrosis factor
TUNEL: terminal deoxynucleotidyl transferase dUTP and nick labeling
TEM: transmission electron microscope
DLS: dynamic light scattering
PALS: Phase Analysis Light Scattering
FBS: fetal bovine serum
MEM: Minimum Essential Medium
VVG: Verhoeff–van Gieson
PCA: Principal Component Analysis
GSEA: Gene Set Enrichment Analysis
miR: microRNA
TCA: tricarboxylic acid
H_2_O_2_: hydrogen peroxide
PKC: protein kinase C
GAP: GTPase-activating protein

## Novelty and Significance

### What is known

- Oxidative stress is implicated in the development and progression of AAA
- Antioxidant therapy has been tried in AAA with no definitive or lasting benefit

### What new information does this article contribute

- Peptide-based delivery of mRNA augments SOD2 expression in aneurysmal tissue
- Enhanced expression of SOD2 mitigates AAA development and largely prevents rupture
- Proteomics confirms that SOD2 overexpression significantly affects oxidative phosphorylation

## Summary

Abdominal aortic aneurysm (AAA) is a condition seen predominantly in men aged 65 years or older. Rupture of AAA carries a mortality rate of greater than 80%. Yet there is currently no selective or molecularly targeted therapy for AAA. Although oxidative stress is implicated in the development and progression of AAA, antioxidant therapy has been unsuccessful due to limitation in delivering sufficient level of antioxidants to the site of AAA. Using a peptide-based nanoplatform, we successfully delivered mRNA to augment the expression level of SOD2 in the aneurysmal tissue to mitigate the progression of AAA and largely prevent rupture. Unbiased proteomic profiling of the aortic tissue in animals administered SOD2 mRNA revealed enrichment in pathways related to inflammation, platelet activity in addition to oxidative phosphorylation, confirming the role of SOD2 in the maintenance of vascular redox balance. The findings suggest that nanoparticle-based SOD2 augmentation offers a promising approach for the non-surgical treatment of AAA. Furthermore, we identify a set of proteins downstream of SOD2 that may be explored as therapeutic targets in AAA. Finally, peptide-based delivery of mRNA can be extended to the modulation of a multitude of pathways underlying human disease processes beyond AAA.

